# The actin-binding protein drebrin interacts with the respiratory syncytial virus matrix protein and promotes virus particle release

**DOI:** 10.64898/2026.10.05.756730

**Authors:** Tristan Jaouen, Katherine Marougka, Manuel Feliciano, Aude Jobart-Malfait, Laila Sago, Monika Bajorek, Marie-Anne Rameix-Welti, Delphine Sitterlin

**Affiliations:** M3P, Institut Pasteur, UMR 1173, Université de Versailles Saint-Quentin, Université Paris-Saclay, Inserm, Université Paris Cité, Paris, France; Unité Virologie et Immunologie Moléculaires, UMR0892, INRAE-UVSQ-Université Paris-Saclay, Jouy-en-Josas, France; Infection & Inflammation, UMR 1173, Inserm, Université de Versailles Saint-Quentin, Université Paris Saclay, Montigny-le-Bretonneux, France; Université Paris-Saclay, CNRS, CEA, Institute for Integrative Biology of the Cell, 91198 Gif-sur-Yvette, France; Centre National de Référence Virus des Infections Respiratoires (CNR VIR), Institut Pasteur, Université Paris Cité, Paris, France

## Abstract

Respiratory syncytial virus (RSV) is an enveloped virus that assembles and buds at the plasma membrane, producing filamentous infectious particles. Unlike many other enveloped viruses, a substantial fraction of RSV virions remains associated with the surface of infected cells, limiting our understanding of the membrane-scission mechanisms responsible for particle release. Here, we identified the actin-binding protein drebrin as a cellular interactor of the RSV matrix (M) protein, both *in vitro* and in infected cells. We demonstrated that RSV infection induces the relocalization of drebrin to actin-rich structures at the cell periphery associated with viral filaments. Using a Transwell-based assay to quantify infectious particles release without mechanically perturbing cells, we confirmed that RSV release is independent of the endosomal sorting complex required for transport (ESCRT) pathway and showed that drebrin depletion reduced RSV release, whereas drebrin overexpression enhanced it. Since drebrin depletion did not significantly affect overall infectious virus production, our results suggest that drebrin may act during the final step leading to membrane scission and RSV particle release. Finally, proteomics analysis identified Transmembrane 9 superfamily member 2 (TM9SF2) as a drebrin-associated protein specifically during infection. TM9SF2 depletion reduced RSV particle release and impaired the RSV-induced relocalization of drebrin to the plasma membrane. Together, these findings identify drebrin and TM9SF2 as host factors regulating RSV release and support a model in which TM9SF2 promotes the recruitment of drebrin to viral budding sites.

**IMPORTANCE:** Respiratory syncytial virus is a major cause of respiratory disease, especially in infants and older adults. To spread, the virus must produce new particles and release them from infected cells, but how this final step occurs remains poorly understood. We identified one cellular protein, drebrin that interacts with the matrix viral protein and helps newly formed virus particles leave infected cells. Drebrin moves to sites where virus particles form and is needed for efficient release of infectious particles. We also found that the cellular protein, transmembrane 9 superfamily member 2, is required for drebrin recruitment to the cell periphery, near the sites of release. Our results suggest that drebrin and transmembrane 9 superfamily member 2 act late in the release process, helping particles separate from the cell surface.

## INTRODUCTION

Human respiratory syncytial virus (RSV) is a major respiratory pathogen and the leading cause of severe lower respiratory tract infections (LRTIs) in young children worldwide. RSV accounts for more than 33 million LTRIs annually in children under 5 years of age, resulting in 3.6 million hospitalizations and 100 000 deaths (1, 2). Although new preventive measures have recently received authorization, including monoclonal antibodies and maternal vaccine (1), no curative treatment is currently available. Consequently, there remains a strong need to develop antiviral molecules targeting specific stages of the viral cycle.

RSV belongs to the order *Mononegavirales*, family *Pneumoviridae*, and genus *Orthopneumovirus*. The viral particle consists of a lipid envelope harboring two major glycoproteins, the fusion (F) protein, which mediates membrane fusion and viral entry and the attachment (G) protein, which is responsible for viral attachment to host cells. Beneath the envelope, the matrix (M) protein forms a layer along its inner surface. Inside the particle, the viral ribonucleoprotein complex (RNP) is composed of the negative-sense genomic RNA encapsidated by the nucleoprotein N, and associated with the viral polymerase complex proteins, consisting of phosphoprotein (P), large polymerase (L), and transcription antiterminator (M2-1) proteins (3, 4).

*In vivo*, RSV infects respiratory ciliated epithelial cells (5), whereas *in vitro* its tropism is broader, encompassing a wide range of cell lines (6). Following release of the vRNPs into the cytoplasm, transcription and replication take place within viral factories formed by liquid-liquid phase separation (LLPS) (7, 8). Newly synthesized vRNPs are then transported to assembly sites via microtubules and by hijacking the endosomal recycling machinery, through a direct interaction between the L protein and Rab11a (9, 10). It is generally accepted that vRNPs assembly and budding occur directly at the plasma membrane, although one study proposed that viral filaments may emerge from recycling endosomes prior to their insertion into the plasma membrane (11). The minimal viral proteins required to orchestrate assembly are P, F and M, as they are sufficient to generate virus-like particles (VLPs) in a minimal transfection system (12). M plays a central role in this process; recent cryo-electron microscopy (cryo-EM) studies have shown that M oligomerizes into a lattice of dimers underlying the plasma membrane (13, 14). In this conformation M presumably interacts with the cytoplasmic tail of F, which has been shown to be required for viral budding (15, 16) and with P to facilitate the recruitment of RNPs to assembly site. Direct binding of P to M dimers requires the N-terminal region and the oligomerization domain of P (17). In addition, the phosphorylation status of M at Thr 205 was shown to affect M oligomerization and is required for the production of infectious virions (18). Following assembly of these components, progeny virions bud from the plasma membrane predominantly as filamentous particles, although a smaller fraction consists of spherical particles (6, 13, 19). Both morphologies have been observed in models of primary airway cell infection (6). The budding step remains poorly understood but unlike most viruses, RSV egress does not result in efficient release of viral particles into the extracellular environment. Indeed, a substantial proportion of these particles remains associated with the surface of infected cells. These, so called cells-associated viral filaments are infectious and favor short-range cell to cell propagation (20), in addition to syncytium-mediated spread driven by the F protein (21). The host factors governing RSV budding and egress remain only partially characterized, although it is well established that the viral cycle is highly dependent on host actin cytoskeletal components. Notably, RSV activates the ARP2/3 complex to promote actin polymerization and formation of actin-rich membrane extensions such as filopodia (20, 22). In addition, cellular GTPases RhoA and Rac1 are required during the late stages of the viral cycle (23–25). One study reported that virus release is independent of Vps4, a key component of the endosomal sorting complex required for transport (ESCRT) machinery that drives membrane scission, while the Rab11-FIP2 protein seems to be critical (26). Nevertheless, the mechanisms underlying the persistent association of viral particle with the plasma membrane remain unclear. In particular, it is unclear whether this phenotype reflects inefficient membrane scission, which may limit the final release of viral particles. In any case, studying RSV release presents substantial experimental challenges. Given the importance of actin-associated proteins in the RSV viral cycle, we focused on drebrin, an actin-binding protein that has been implicated in filopodia formation and scission process (27, 28).

Drebrin E (Development-Regulated Brain Protein) is 649-amino-acid actin-binding protein that was first reported to be localized in the neuronal growth cone during development (29, 30) and subsequently in the cytoplasm of various cell types including respiratory cells (31, 32). Drebrin is involved in a variety of cellular processes including neurite outgrowth (29) and cell migration, particularly in neurons (33, 34). More broadly, drebrin promotes the formation and stabilization of filopodia in both neuronal and non-neuronal cell lines (27, 29, 35). Drebrin’s functions are mediated by its interaction with F-actin. It binds F-actin through two alpha-helical domains, which were recently characterized by cryo-EM (36). Through its direct interaction with F-actin, drebrin inhibits actin depolymerization, slows barbed-end growth and promotes bundling of actin filaments (37, 38). In addition, drebrin competitively displaces other actin-binding protein (ABPs) from F-actin, such as F-actin disassembly-promoting factor cofilin (39). Drebrin also contributes to the coupling of actin filaments to microtubules through its interaction with the plus-end tracking protein EB3 (29). Moreover, drebrin has also been implicated in the formation of intracellular vesicles during endocytosis (40, 41) and in the production of extracellular vesicles through ectocytosis at the primary cilium of cells (28). These observations raise the possibility that drebrin may also participate in membrane remodeling and scission processes.

In this study, we performed affinity purification of the M protein and identified drebrin as a key interactor of M in infected cells. We then demonstrate, using an *in vitro* assay that this binding is direct. We set up a novel assay to quantify viral particle release and we show that drebrin is required for this step. This result is consistent with our observation that RSV infection relocalizes drebrin to the cell periphery, where it accumulates at virus-induced filopodia and viral filaments, suggesting that drebrin is recruited to viral assembly and egress sites. Finally, we identify TM9SF2 as an infection-specific interactor of drebrin and show that TM9SF2 is also required for efficient RSV particle release and contributes to the relocalization of drebrin to the cell periphery during infection. Altogether, our results provide new insights into the mechanisms governing budding and egress of the RSV.

## RESULTS

### The RSV M protein interacts directly with the cellular protein drebrin *in vitro* and in infected cells

To investigate the role of the drebrin protein in RSV release, we first assessed its interaction with the RSV M protein, as viral matrix proteins frequently play central roles in membrane scission. HEp-2 cells were infected with RSV for 24 h and whole-cell lysates were subjected to immunoprecipitation (IP) using anti-M antibodies. Total lysates, unbound and bound fractions were then analyzed by Western blotting for M, actin and drebrin proteins. As shown in Fig. 1A, both drebrin and actin were detected in the M-bound fraction, suggesting an interaction between M, actin and drebrin in infected cells. To confirm this interaction, we performed a reverse IP targeting drebrin. HEp-2 cells stably harboring a replicating episomal plasmid expressing either YFP-tagged drebrin (drebrin-YFP) or YFP alone as a negative control, were infected with RSV. At 24 h post-infection, cell lysates were incubated with agarose beads covalently coupled to an anti-GFP nanobody (VHH) to immunoprecipitate YFP and drebrin-YFP proteins. Bound proteins were then analyzed by Western blotting using antibodies against M and YFP. As shown in Fig. 1B, M was specifically detected in the bound fraction from infected cells expressing drebrin-YFP, whereas no signal was detected in cells expressing YFP alone. To assess whether this interaction also occurs in non-infectious condition, we performed a drebrin IP in HEp-2 cells transiently expressing M protein. M was detected in the drebrin IP under these conditions as well (Fig. S1). Taken together, these results support an interaction between drebrin and the RSV M protein in both infected and transfected cells.

**FIG 1.**
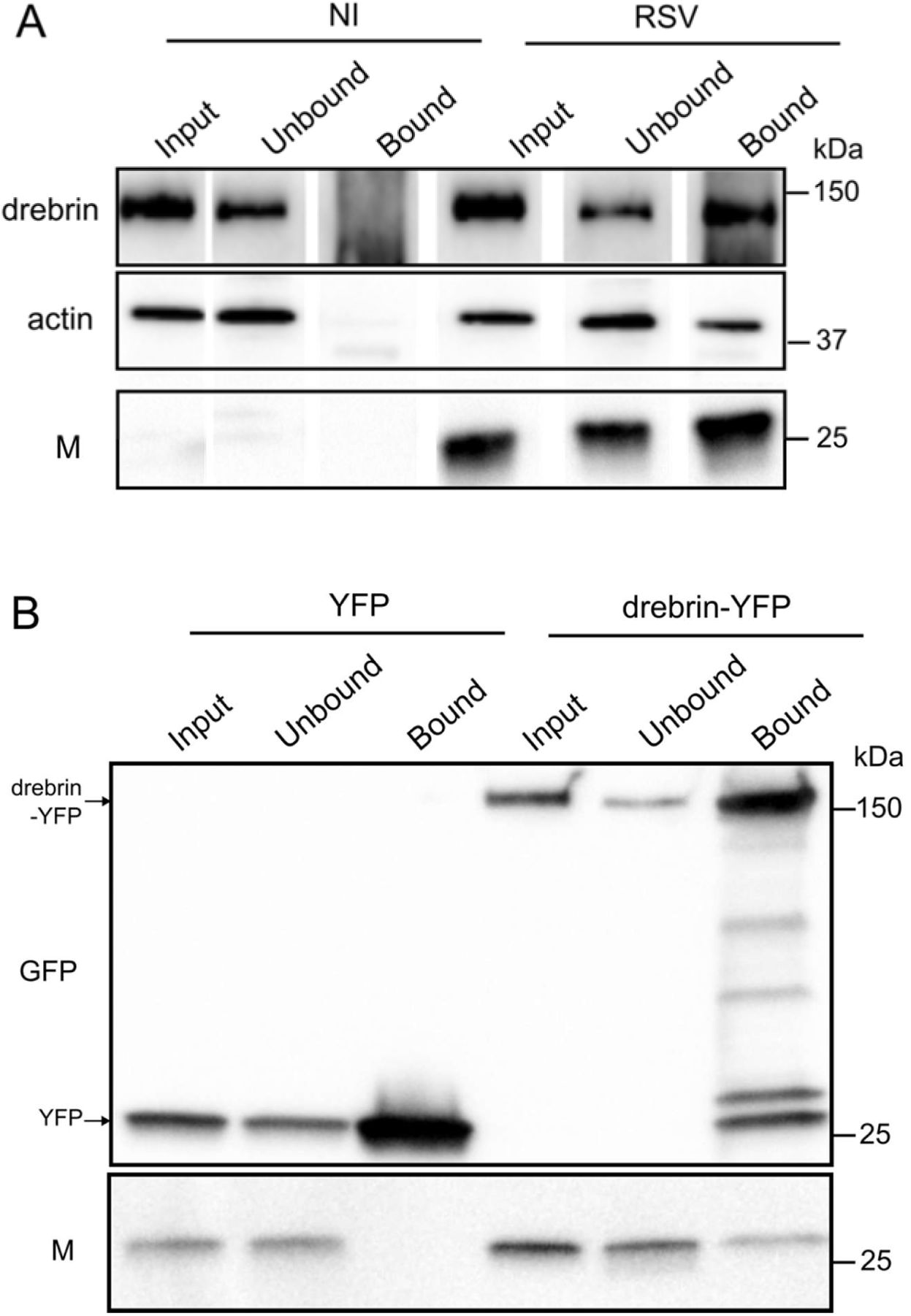
Drebrin interacts with M in infected cells. (A) HEp-2 cells were “mock” infected (NI) or infected with RSV for 24 h, at MOI 1 and processed for IP using M antibodies. Input (1 %), unbound (1 %) and bound (30 %) fractions were loaded on SDS-PAGE and a Western blot was performed using antibodies indicated on the left. Representative of three independent experiments. (B) HEp-2-Drebrin-YFP and HEp-2-YFP cells were infected with RSV for 24 h, at MOI 1 and processed for IP using anti-GFP nanobodies. Input (0,5 %), unbound (0,5 %) and bound (30 %) fractions were loaded on SDS-PAGE and a Western blot was performed using antibodies indicated on the left. Drebrin-YFP and YFP corresponding bands were indicated by an arrow on the left. Representative of three independent experiments.

We next investigated whether the M-drebrin interaction occurs through direct binding. To address this question, recombinant proteins were produced and purified *in vitro.* M has previously been shown to form dimers upon purification (42), whereas drebrin was most likely purified as oligomers (Fig. S2). The potential interaction between the two proteins was then assessed by native agarose gel electrophoresis. As shown in Fig. 2A increasing concentrations of drebrin protein progressively shifted the migration profile of M on native agarose gel, demonstrating a direct protein-protein interaction. To assess the specificity of this interaction, the RSV P was used as negative control. Because wild-type (WT) P exhibited a migration profil similar to that of drebrin, we also tested a series of P truncation mutants. As shown in Fig. 2B, neither the WT or the P mutants altered the migration profile of drebrin, supporting the specificity of the interaction detected between drebrin and M.

**FIG 2.**
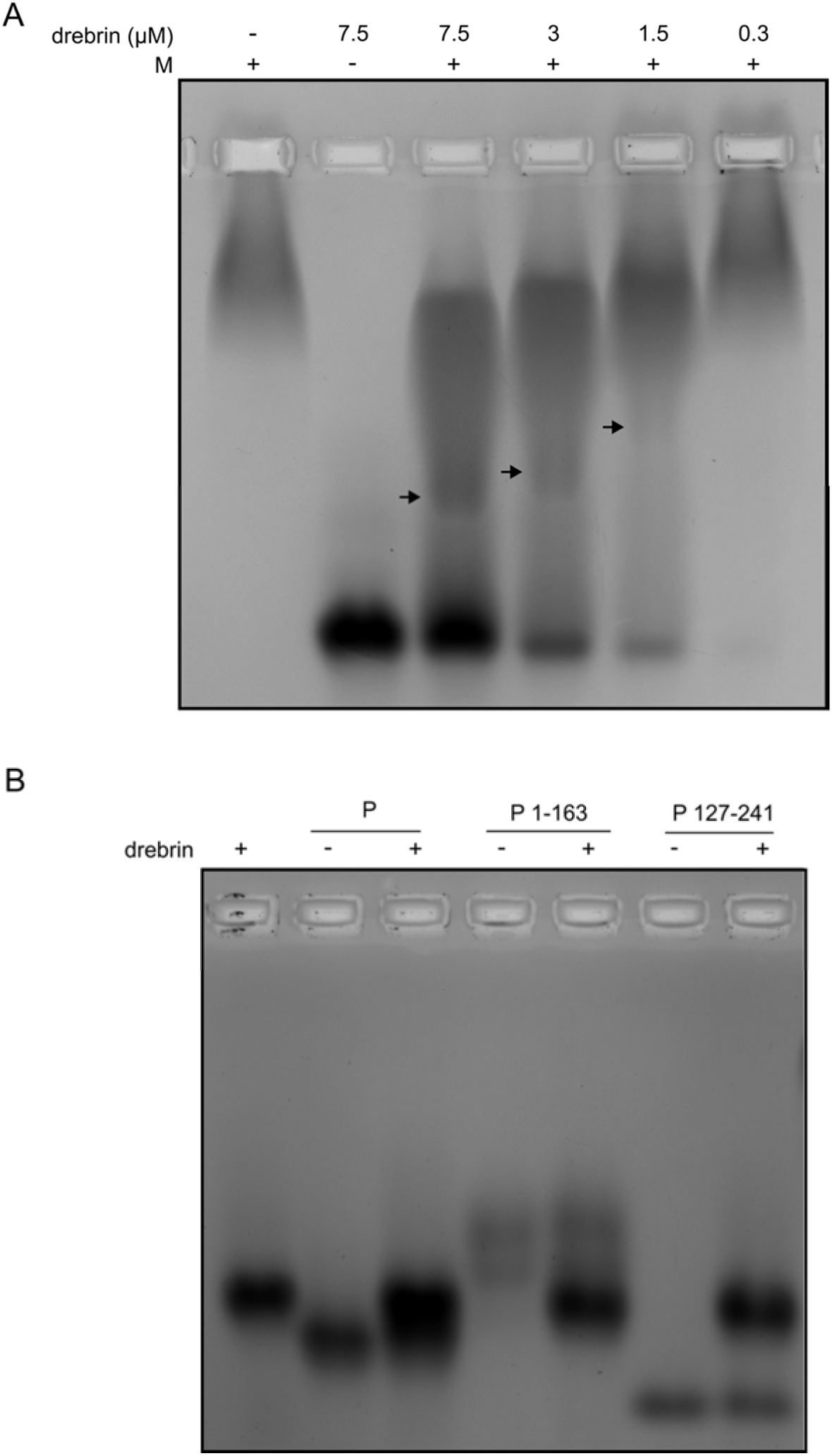
**Drebrin interacts with M *in vitro.*** (A) purified M (15 µM) and increasing amounts of drebrin or (B) purified drebrin (7.5 µM) and P or P deletion constructs (15 µM) were incubated 20 min RT prior to analysis of formation of complexes by band shift on native agarose gel. Arrows indicate complex formation. Each image is representative of three independent experiments.

### RSV infection drives drebrin relocalization to the cell periphery at the viral budding sites

We next assessed the expression of drebrin and its subcellular localization in RSV- infected cells at the end of the viral cycle (24 h post-infection). First drebrin mRNA level were examined in infected and non-infected cells by RT-qPCR analysis on whole cell lysates. As shown in Fig. 3A drebrin mRNA level is not significantly altered during infection. Consistent with this finding, Western blot analysis showed that the total level of drebrin protein remained unchanged in infected cells compared to non-infected cells (Fig. 3B and 3C). We then explored the subcellular localization of the endogenous drebrin protein by immunofluorescence in both non-infected and infected cells. As shown in Fig. 3D, drebrin localization differed remarkably between the two conditions, shifting from a predominantly cytoplasmic distribution in non-infected cells to a nearly plasma membrane localization in RSV-infected cells (broad field view in Fig. S3). We quantified the ratio of the mean fluorescent intensity at the cell periphery to that in the intracellular cytoplasmic compartment. This analysis confirmed a significant increase in cell periphery-associated drebrin in infected cells (Fig. 3E). Overall, these results show that RSV infection does not affect drebrin expression levels but rather induces its relocalization toward the plasma membrane.

**FIG 3.**
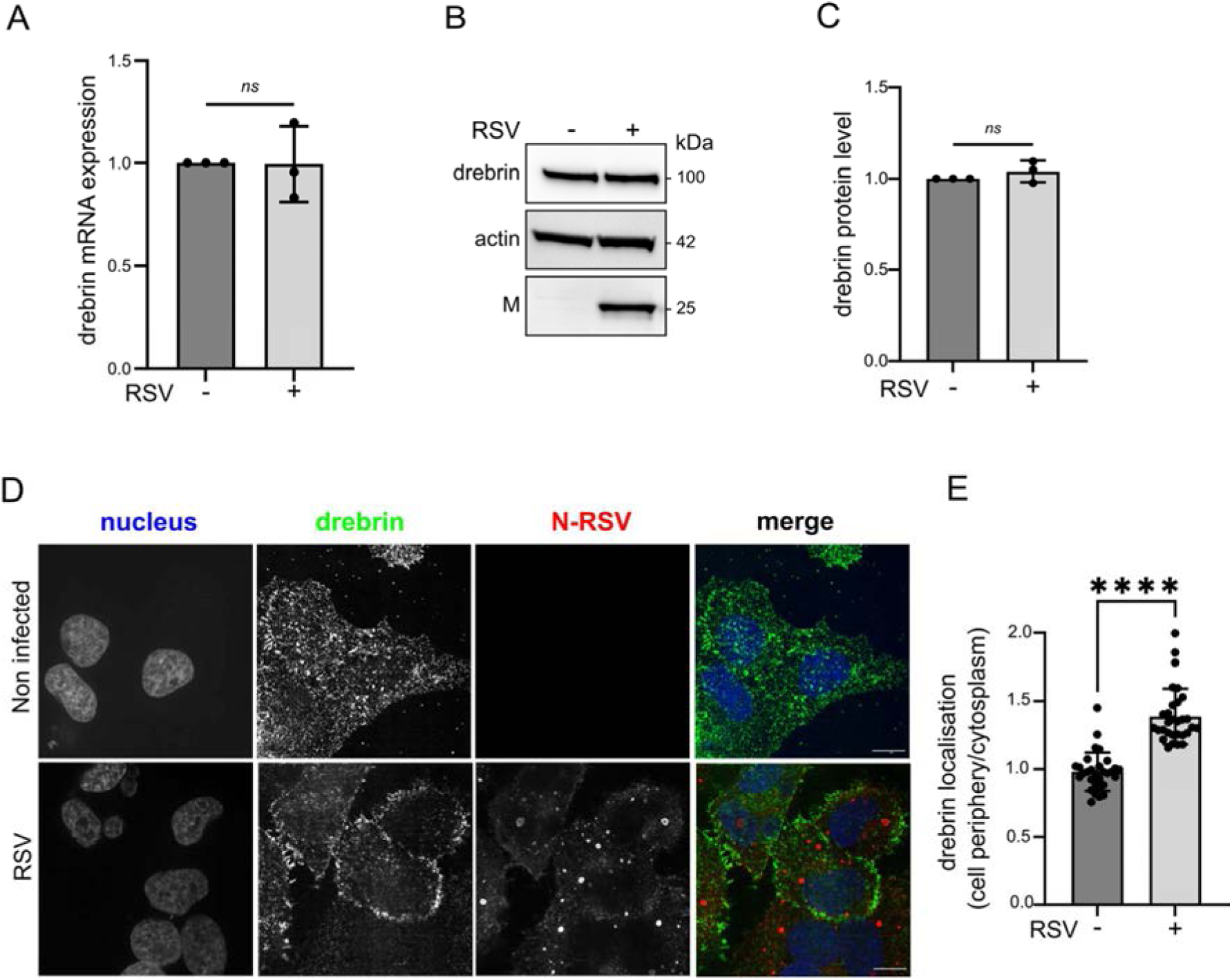
Drebrin is localized at cell periphery in infected cells. HEp–2 cells were infected or not with RSV for 24 h at MOI 1. (A) Drebrin mRNA level was quantified by RT-qPCR. Data were analyzed using the 2 ^- ΔΔCt^ method. GAPDH mRNA was used to normalize the mRNA quantity between conditions. Bar graph shows mean ± standard deviation (SD) from three independent experiments. One sample t test was performed *ns* : *P* > 0.05. (B) Total protein extracts were analyzed by Western blot with antibodies indicated on the left. (C) Quantification of Drebrin protein signal (shown in B) was performed using the ImageLab Software as described in methods section. Bar graph shows mean ± SD from three independent experiments. One sample t test was performed *ns* : *P* > 0.05. (D) Cells were stained with antibody against drebrin (green in merge) and N-RSV (red in merge). Hoechst staining is shown in blue in merge. Representative images from 3 independent experiments are shown. One representative Z step is shown. Scale bar 10 μm. (E) Quantification of drebrin signal (shown in D) was performed using the ImageJ Software as described in methods section. The drebrin mean fluorescence intensity at the cell periphery was normalized to the drebrin mean fluorescence intensity in the cytoplasmic. Bar graph shows mean ± SD from 2 independent experiments (15 cells per condition per experiment were analyzed). Mann-Whitney test was performed, *\*\*\*\** : *P* < 0.0001.

These results prompted us to investigate the localization of drebrin relative to viral budding sites. To address this question, immunofluorescence analysis was performed to simultaneously visualize actin, drebrin and viral particles using an antibody directed against the RSV envelope G protein. As shown in Fig. 4A, a clear co-staining between actin and drebrin was observed in RSV infected cell, particularly at filopodia-like structures that form during infection. Importantly, higher magnification view revealed that drebrin-enriched membrane domains correspond to site of viral particle emergence, suggesting that drebrin accumulates specifically at viral budding sites. As M interacts with drebrin, we next examined the subcellular localization of both proteins using expansion microscopy. As shown in Fig. 4B, viral filaments were readily detected by M antibody staining, whereas drebrin labeling was weaker, with no clear evidence of its relocalization to the cell periphery, likely due to reduced antibody recognition in expanded gels. Nevertheless, punctate drebrin signals were frequently observed along M-positive viral filaments. Collectively, these results demonstrate that drebrin is recruited at viral budding sites and suggest that drebrin may be retained with viral filament in association with the matrix protein.

**FIG 4.**
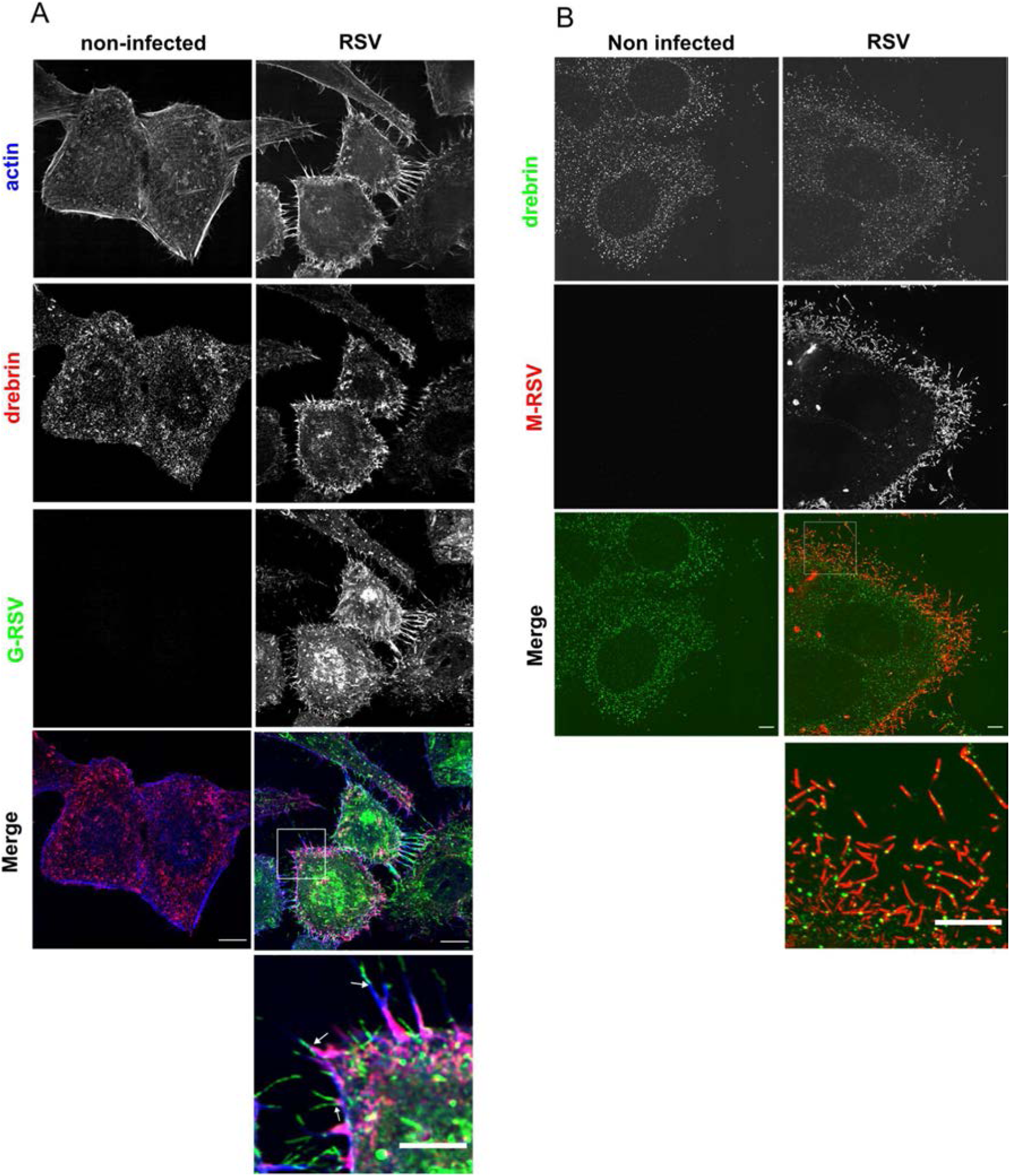
**Drebrin is localized at the budding site of viral particle**. HEp–2 cells were infected or not with RSV for 24 h at MOI 1. (A) Cells were stained with an antibody against drebrin (red in merge) and G-RSV (green in merge) and with phalloidin (blue in merge). Representative images from 3 independent experiments are shown. One representative Z step is shown. Scale bar 10 μm. Zoom image of the white square are shown below the merge (scale bar 5 µm). White arrows indicate drebrin and actin-enriched membrane regions from which viral particles emerge. (B) Cells were processed for expansion microscopy and stained with an antibody against drebrin (green in merge) and M-RSV (red in merge). Representative images from 3 independent experiments are shown. Projection of 8 Z step is shown. Zoom image of the white square are shown below the merge. Scale bar 5 µm.

### Drebrin promotes RSV viral particle release

We next examined whether drebrin depletion affected the total production of infectious RSV virus particles over a single replication cycle. At 24 h pi, total virus production, including both cell-free and cell-associated virions, was quantified by plaque assay, in cells treated with drebrin-specific siRNA or non-targeting (NT) siRNA. In parallel, drebrin depletion was confirmed by Western blotting 2 days after siRNA transfection (Fig. S4A-B). No cytotoxicity was detected under these conditions (Fig. S4C). As shown in Fig. 5A no significant difference in total infectious virus production was observed between drebrin-depleted and control cells. Viral multiplication was also monitored by measuring mCherry fluorescence over 24 h following infection with a recombinant RSV-mCherry in the presence or absence of drebrin. The kinetics of mCherry signal were comparable between the two conditions and fluorescence intensity at 24 h post-transfection did not differ significantly from that measured in NT siRNA-treated cells (Fig. 5B and 5C). Together, these results indicate that drebrin does not affect the overall production of infectious RSV particles. As drebrin is relocalized to the cell periphery during the late stages of the viral cycle, we next examined whether its depletion affected the formation of viral filaments. HEp-2 cells treated with drebrin-specific or NT siRNA were infected with RSV for 24 h and immunofluorescence were performed to visualize, drebrin and viral particles using antibody against G. As shown in Fig 5D, no apparent qualitative difference in viral filament formation was observed between drebrin-depleted and control cells.

**FIG 5.**
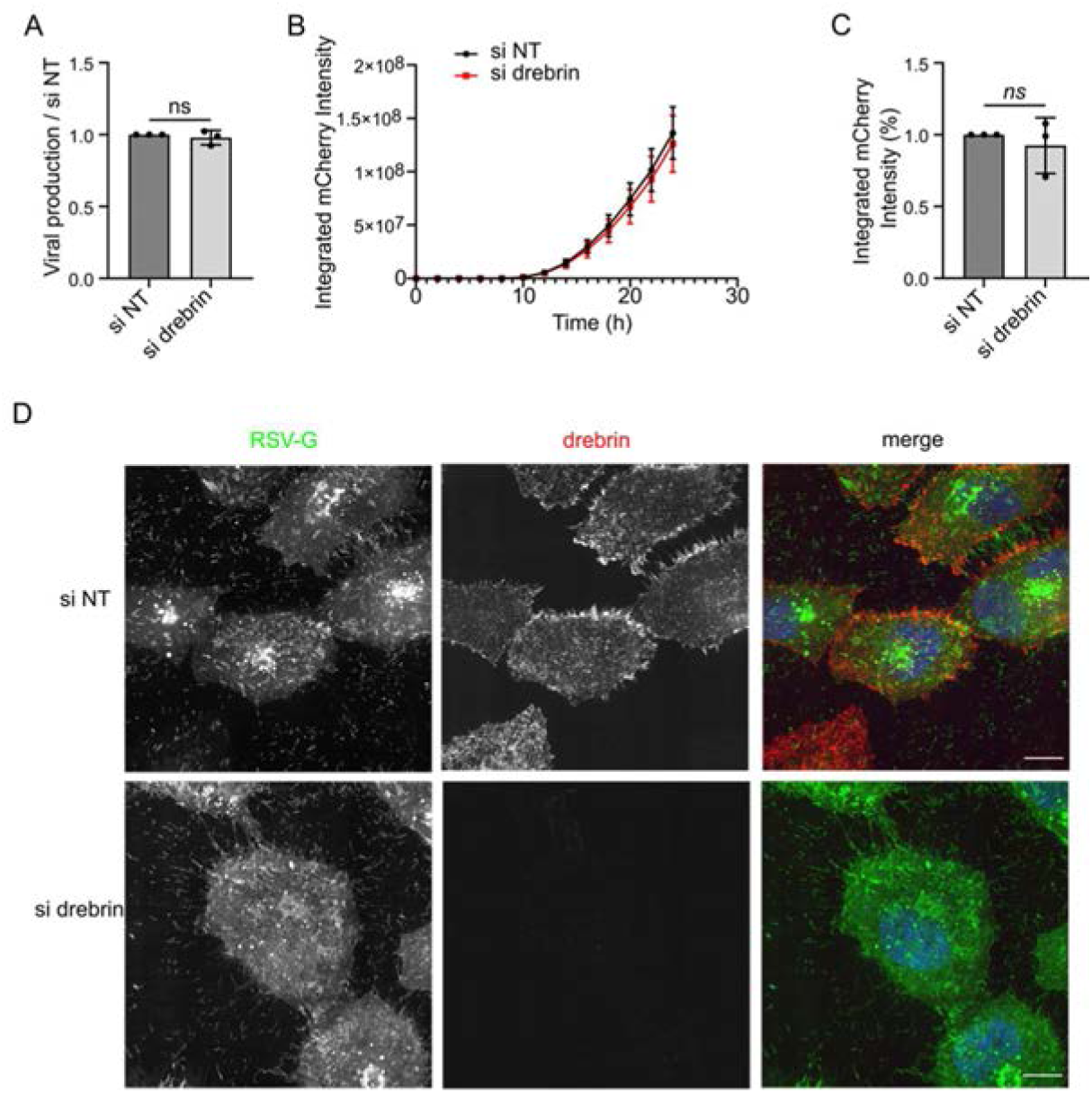
Drebrin does not affect production of infectious RSV particles. (A) HEp-2 cells were transfected with control (si NT) or drebrin siRNAs (si drebrin) for 48h and subsequently infected with RSV-WT (A,D) or RSV-mCherry (B-C) at MOI 1 for 24 h. (A) Viral titers were determined by plaque assay and normalized to the si NT condition. Bar graph shows mean ± SD from 3 independent experiments performed in triplicate. A one-sample t-test was performed. *ns* : *P* > 0.05. (B) Graph of mCherry signal at different time post-infection in si NT (black) and si drebrin (red) infected cells. The data shown are the mean ± SD of 3 independent experiments performed in triplicate. (C) mCherry intensity in si drebrin cells at 24 h post-infection relative to si NT. A one sample t-test was performed, *ns* : *P* > 0.05. (D) Cells were stained with an antibody against drebrin (red in merge) and G-RSV (green in merge) and with phalloidin (blue in merge). Representative images from 3 independent experiments are shown. One representative Z step is shown. Scale bar 10 μm.

A previous study has implicated drebrin in ectocytosis (28), we therefore investigated whether drebrin contributes to RSV particle release. As described in the introduction, RSV release from HEp-2 cells is incomplete, with the majority of RSV particles remaining loosely associated with the cell surface. In our experience, conventional collection of the culture supernatant provided a poorly reproducible estimate of the cell-free RSV fraction, potentially because this fraction is very sensitive to mechanical perturbations caused by pipetting. We therefore developed a rapid and sensitive assay to quantify the release of RSV particles into the culture supernatant without any mechanical disruption of the infected cells. In this assay HEp-2 cells were seeded in the upper compartment of a 0.4 µm pore size transwell insert, and infected with RSV-mCherry virus. Following one complete viral multiplication cycle, neosynthesized viral particle released from the donor cells diffused through the porous membrane of the transwell and infected target cells seeded in the lower chamber (Fig. 6A). Cell infection was monitored by measuring mCherry fluorescence intensity using an IncuCyte live-cell imaging system. The fluorescence signal in the target-cell population, normalized to that in the donor-cell population, was used as a quantitative readout of viral particle release efficiency. This assay was first validated using as a positive control, cells depleted for a subunit of Arp2/3 complex, known to be required for release (Fig. S5). We also confirmed that RSV release is ESCRT independent, using cells depleted for CHMP4B, an ESCRT-III complex component, and for VPS4, the AAA+ ATPase required for the final disassembly of ESCRT-III complex (Fig. S5). Next, we addressed the role of drebrin in viral release using cells depleted for drebrin. As shown in Fig. 6B drebrin depletion significantly reduced the amount of viral particle released into the supernatant of infected cells. Conversely, performing the same assay in cells overexpressing drebrin-YFP revealed a significant increase in viral particle release compared with cells overexpressing YFP (Fig. 6C). As drebrin depletion did not affect total viral production, these findings indicate that drebrin specifically contributes to the release of viral particles. The absence of a detectable effect in the total virus production is consistent with the fact that the released fraction represents only a minor proportion of the total viral production. Collectively, these results suggest a role of drebrin in the final membrane scission step that governs viral particles release.

**FIG 6.**
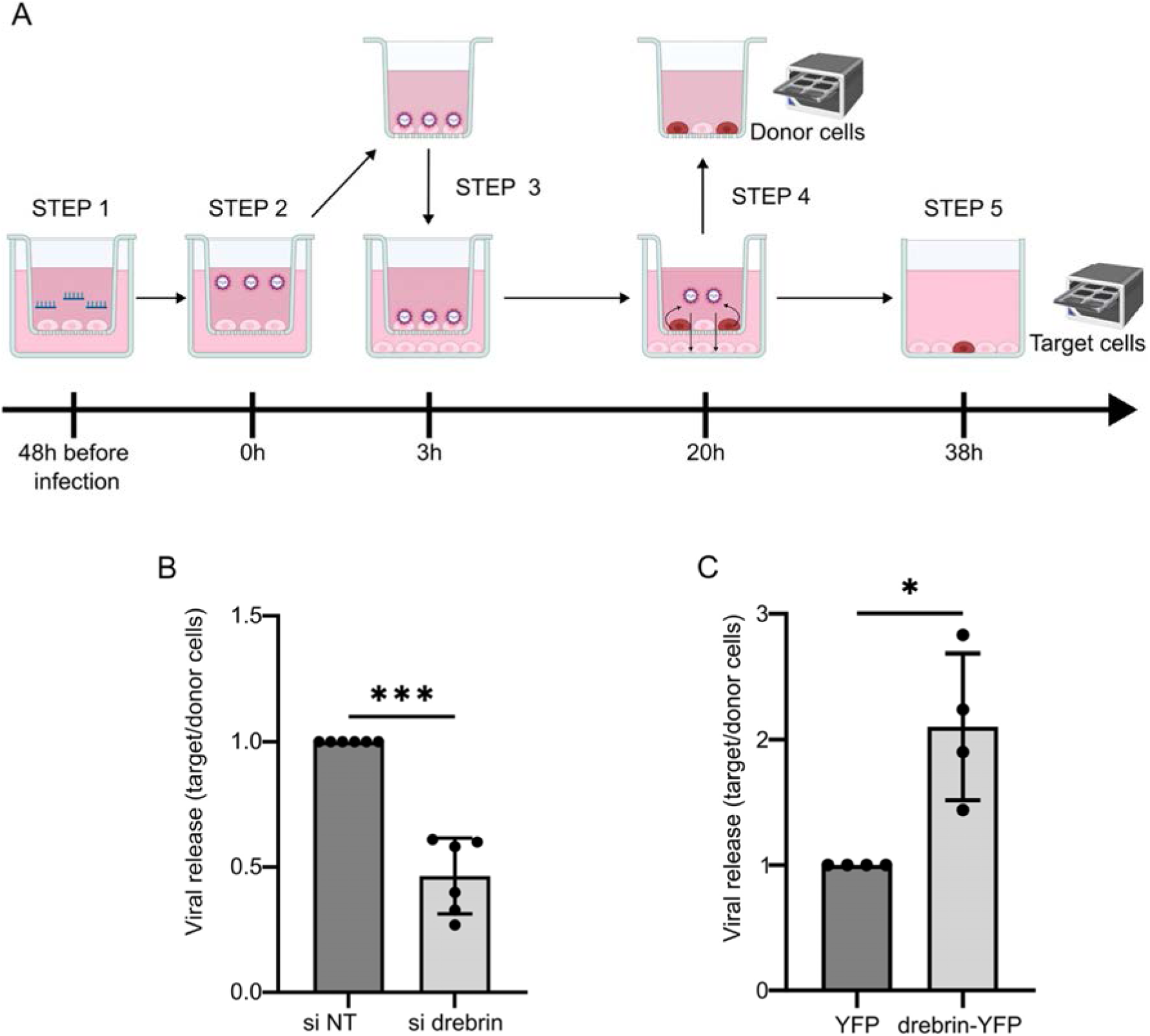
**Drebrin promotes viral particle release**. (A) Schematic representation of the release assay. HEp-2 cells were transfected with control or specific siRNAs and seeded onto Transwell inserts for 48h (step 1). Cells were then infected with RSV-mCherry at an MOI of 1 (step 2). After washes to remove non-internalized virus, the infected “donor” cells were then placed in contact with target cells (step 3). The number of infected “donor” and “target” cells were quantified based on mCherry fluorescence using the Incucyte S3 system at 20h (step4) and 38h (step 5) post-infection respectively. RSV release efficiency was calculated as the ratio of infected target cells to infected donor cells and normalized to the control siRNA condition. (B) HEp-2 cells were transfected with si NT or drebrin-specific siRNAs (si drebrin) and release assay was performed as described in A. Date are mean ± SD of 6 independent experiments, each point represents mean of triplicates. Y axis represents viral release efficiency (ratio of infected target cells to infected donor cells) normalized to the control siRNA condition. (C) release assay was performed with HEp-2-YFP and HEp-2-Drebrin-YFP cells. Data are mean ± SD of 4 independent experiments done in triplicates. Y axis represents viral release efficiency (ratio of infected target cells to infected donor cells) normalized to the control siRNA condition. (B-C) A one-sample t-test was performed. *\** : *P* < 0.05; *\*\** : *P* < 0.001.

### The actin binding and intrinsically disordered domains of drebrin are required for M interaction

To better understand the mechanism by which drebrin contributes to RSV particle release, we first mapped the drebrin domain required for interaction with the viral M protein. Drebrin comprises an ADFH domain, two actin-binding domains (ABD1 and ABD2) and a large C-terminal intrinsically disordered domain (IDD) (Fig 7A). Hep-2 cells stably expressing four drebrin-YFP truncation mutants were generated and stably expressed in HEp-2 cells using the episomal plasmid-based system described above. The interaction between the drebrin mutants and M was then analyzed by IP in RSV-infected cells. M coimmunoprecipitated with the WT drebrin and with the ABD1-ABD2-IDD mutant, whereas no interaction was detected when the ABD1-ABD2 and IDD regions were expressed separately (Fig. 7B and S6). Analysis of the input fractions showed comparable expression levels for most constructs, with the exception of the ABD1-ABD2-IDD mutant, which appeared to be expressed at a lower level. These results indicate that efficient association between drebrin and M requires the combined presence of the ABD and IDD domains. In parallel, the subcellular localization of the mutants was examined. Consistent with previous observations (27), the presence of the ABD regions was required for the filopodia-like localization of drebrin (Fig. S7). We next assessed the functional impact of drebrin mutants overexpression on RSV particle release, using the assay described in Fig. 6A. As expected, overexpression of WT drebrin-YFP enhanced RSV viral particle release. In contrast, overexpression of the ADFH-ABD1-ABD2 mutant, which did not interact with M, failed to promote particle release and instead significantly reduced it (Fig 7C). Overexpression of the ABD1-ABD2-IDD mutant, which displayed a stronger interaction with M, had a detectable but weaker effect on viral release than WT drebrin (Fig 7C). Taken together, these results suggest that the drebrin domains required for M interaction retain the ability to promote RSV release.

**FIG 7.**
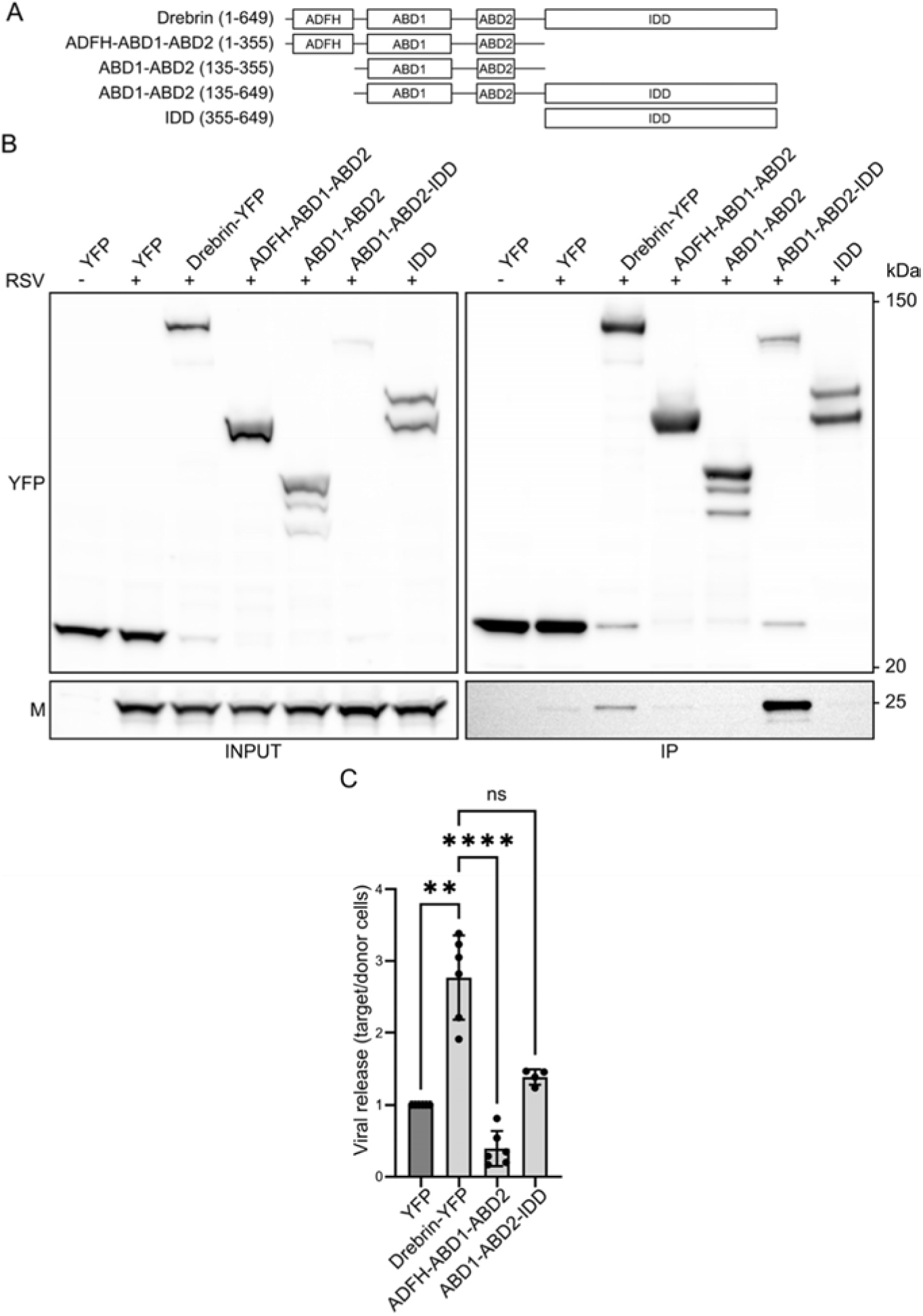
ABD and IDD domains of drebrin are required for M interaction but not for viral particle release. (A) diagrams of drebrin mutants are shown. ADFH, actin-depolymerizing factor homology domain. ABD1, actin binding domain 1. ABD2, actin binding domain 2. IDD intrinsically disordered domain. Constructs are YFP-tagged at the C-terminus. Numbering is for human drebrin E. (B) HEp-2 cells lines were established with the indicated plasmids and cells were infected with RSV for 24h at MOI 1 and processed for IP using anti-GFP nanobodies. Input (0,5 %), unbound (0,5 %) and bound (30 %) fractions were loaded on SDS-PAGE and Western blotting using antibodies indicated on the left. Representative of three independent experiments. (C) release assay was performed as in Fig 6C with cells expressing indicated plasmids. Data are mean ± SD of 6 independent experiments done in triplicates. Y axis represents viral release efficiency (ratio of infected target cells to infected donor cells) normalized to the control siRNA condition. One-Way ANOVA with Dunnett’s multiple comparisons test and Geisser-Greenhouse correction was performed, *ns* : *P* > 0.05, *\*\** : *P* < 0.01, *\*\*\*\** : *P* < 0.0001.

### The infection-specific drebrin interactome reveals TM9SF2 as a positive regulator of RSV particle release

Our results showed that drebrin relocalizes to the plasma membrane during RSV infection (Fig. 3). We therefore hypothesized that this relocalization might be accompanied by changes in its protein interaction network. Characterization of the infection-specific drebrin interactome could thus reveal novel cellular factors involved in RSV particle release. To this end, we used the HEp-2 drebrin-YFP cell line and the immunoprecipitation protocol established above to characterize drebrin-associated proteins at 24 h post-infection relative to non-infected controls. Drebrin-YFP associated proteins were identified by mass spectrometry and comparative analysis revealed 26 cellular proteins that were significantly enriched in RSV infected HEp-2-drebrin-YFP cells compared with non-infected cells (Fig 8A and Table S1). Note that vRNPs proteins, as well as the F and M viral proteins were also identified. However, their specific enrichment cannot be assessed in this analysis, since comparisons were made using non-infected control cells. To identify cellular proteins potentially involved in RSV release, we first excluded proteins corresponding to interferon-stimulated genes whose upregulation during infection, could suggest that their enrichment in the drebrin interactome under infection conditions might be an artefact. We then used RT-qPCR to compare the mRNA expression levels of the remaining 17 candidate proteins in infected and non-infected cells. Eight candidates displayed more than 3-fold increase in mRNA levels following infection (Fig. 8B and Fig. S8A), suggesting also that their enrichment in the drebrin interactome under infection conditions might be an artefact. We next assessed the effects of candidate depletion on viral replication and excluded 3 more candidates whose depletion modulated viral replication more than 40 % (Fig. 8B and Fig. S8B). We focused on six candidates with no or limited effect on replication, allowing us to specifically assess their contribution to viral release. The contribution of these candidates to RSV particle release was analyzed using the viral release assay (Fig. 6A). HEp-2 cells were transfected with siRNAs targeting TM9SF2, CTNNA1, CTNNA2, ANXA5, FUBP3 or PGLS, and viral release was quantified as described above. In parallel, siRNA-mediated depletion and cytotoxicity were controlled 48 h post-transfection (Fig. S9). As shown in Fig. 8C, depletion of 5 candidates had no significant effect on viral release, although depletion of CTNNA1 and 2 showed a tendancy for increased viral particle release, suggesting that these proteins could negatively regulate this process. Notably, depletion of TM9SF2 strongly reduced viral particle release, identifying TM9SF2 as a potential positive regulator of RSV release. To determine whether TM9SF2 affects total virus production, HEp-2 cells, transfected with control or TM9SF2-specific siRNAs, were infected with RSV, and infectious virus production was quantified 24h p.i. by plaque assay (Fig. S10). TM9SF2 depletion did not significantly affect total viral production compared with control cells indicating that, like drebrin, TM9SF2 specifically promotes the efficient release of RSV particles.

**FIG 8:**
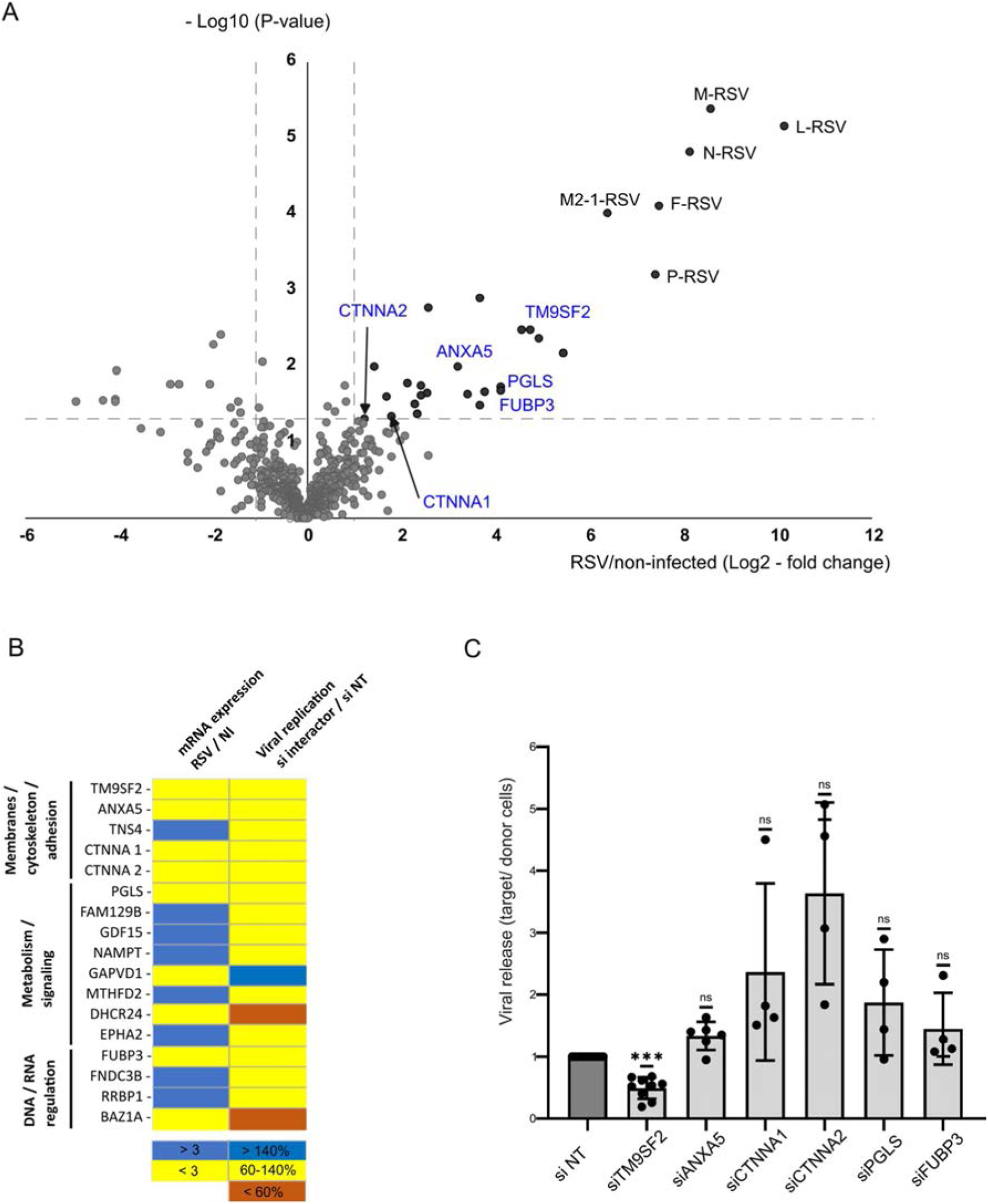
TM9SF2 is a drebrin interactor, required for RSV viral release. (A) Drebrin-YFP HEp-2 cells were uninfected or infected with RSV at MOI 1 for 24 h. Proteins associated with drebrin-YFP were affinity-purified using GFP nanobody-coupled agarose beads, identified by mass spectrometry, and quantified based on spectral counts obtained for each sample. A log2 fold change was calculated by comparing RSV-infected and uninfected conditions. Only proteins with at least two spectral counts in at least two biological replicates in either condition were included in the analysis. Statistical significance was assessed using a beta-binomial test adapted for spectral count data. Results are presented as a volcano plot. The horizontal dashed gray line indicates the statistical significance threshold (*P* < 0.05), whereas the vertical dashed gray lines indicate a twofold difference between infected and uninfected samples. Cellular proteins used in C are indicated in blue. Viral RSV proteins are indicated in black. Results from three independent biological replicates (B) Left: Heat map of RT-qPCR analysis comparing mRNA levels of 17 candidate proteins in RSV-infected and uninfected cells. Blue indicates > 3-fold upregulation and yellow < 3-fold. Right: Heat map of viral replication analysis comparing RSV-mCherry signal in RSV-infected cells treated with interactors-specific siRNA and si NT. Blue indicates replication >140%, brown replication < 60% and yellow 60% < replication >140%. Complete data are shown in Fig S8. (C) HEp-2 cells were depleted with the indicated siRNAs and viral release assay was performed and compared with control conditions. One-Way ANOVA with Dunnett’s multiple comparisons test and Geisser-Greenhouse correction was performed relative to si NT. *\*\*\** : *P* <0.001, *ns* : *P* >0,05.

### TM9SF2 is required for the RSV-induced relocalization of drebrin to the cell periphery

TM9SF2 is a transmembrane protein localized to endosomal and trans-Golgi network membranes. Although its function remains poorly characterized, TM9SF2 has been implicated in glycosylation processes (43, 44). Since suitable antibodies were unavailable and overexpression of epitope-tagged TM9SF2 was cytotoxic, we used a loss-of-function approach to investigate more precisely its role during RSV infection. To determine whether TM9SF2 depletion affects the localization of endogenous drebrin, HEp-2 cells were transfected with NT or TM9SF2-specific siRNAs and infected or not with RSV for 24 h, before IF analysis. As observed previously, RSV infection induced a pronounced redistribution of drebrin toward the plasma membrane in NT siRNA-treated cells (Fig. 9A). Interestingly, this infection-induced relocalization was strongly impaired following TM9SF2 depletion. Quantitative analysis shows that TM9SF2 depletion significantly reduced the RSV-induced relocalization of drebrin to the cell periphery in RSV infected cells (Fig. 9B).

**FIG 9:**
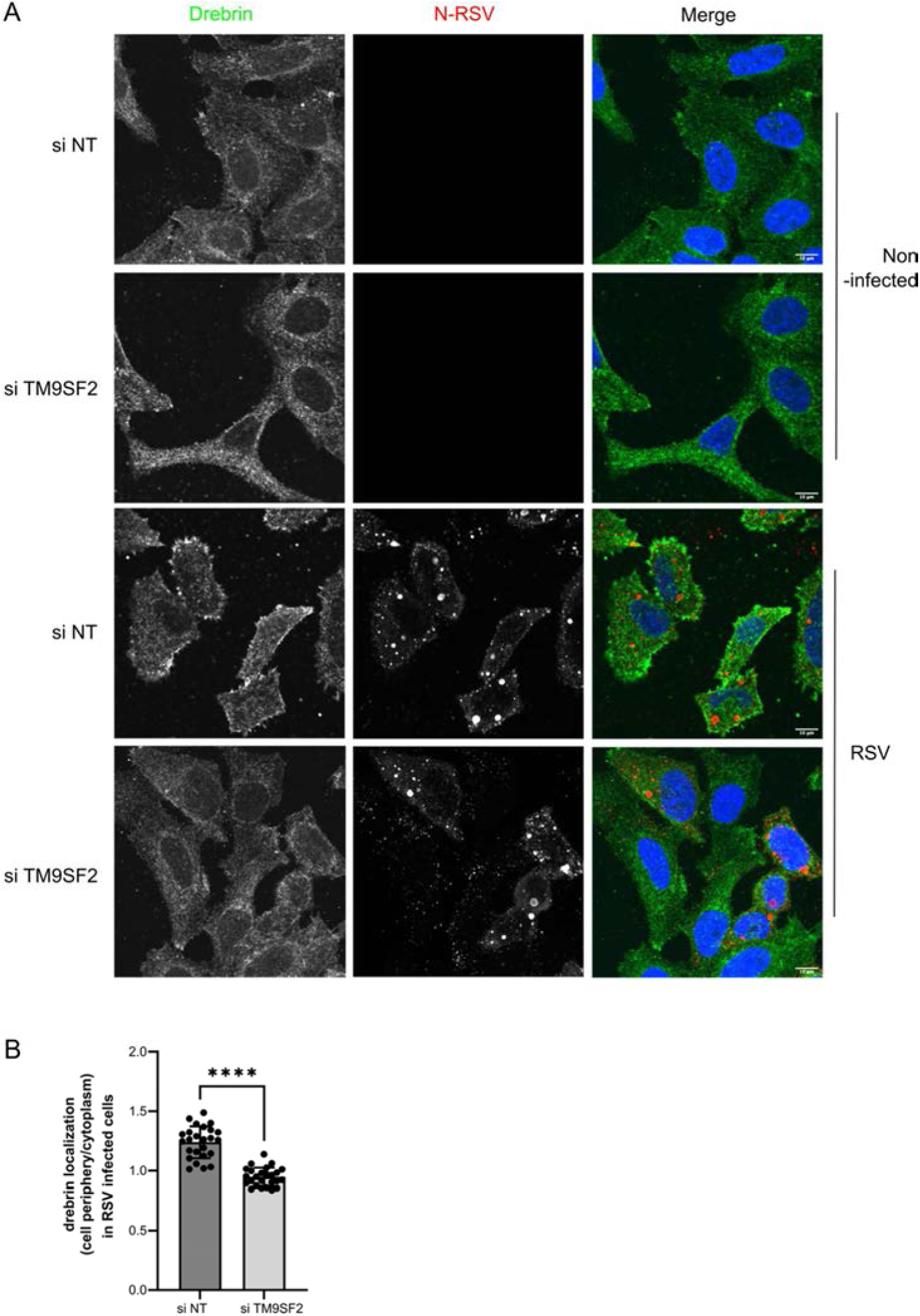
TM9SF2 is required for the Drebrin relocalization at the plasma membrane upon RSV infection. (A) HEp–2 cells were infected or not with RSV-WT at MOI 1. At 24h p.i. cells were fixed and stained with an antibody against Drebrin and N antibodies. Hoechst 33342 staining is shown in blue. Representative images from 3 independent experiments are shown. One representative Z step is shown. Scale bar 10 μm. (B) Quantification of Drebrin signal in RSV infected cells was performed using the ImageJ Software as described in methods section. The drebrin mean fluorescence intensity at the plasma membrane was normalized to the drebrin mean fluorescence intensity in the cytoplasmic. Bar graph shows mean ± SD from 2 independent experiments (15 cells per condition per experiment were analyzed). Welch’s t-tests was performed, *\*\*\*\** : *P* < 0.0001.

Together, these results indicate that TM9SF2 is required both for the RSV-induced redistribution of drebrin to the plasma membrane and for efficient viral particle release. These findings are consistent with a model in which drebrin and TM9SF2 function within a common molecular pathway that promotes RSV particle release, with TM9SF2 acting upstream of drebrin to facilitate its recruitment at the viral assembly and budding sites.

## DISCUSSION

The membrane scission mechanism that allows the release of RSV particles from infected cells into the extracellular environment remains poorly characterized. This process is particularly difficult to investigate because RSV filaments are predominantly retained at the plasma membrane and are therefore not readily recovered from culture supernatants. The development of a quantitative assay to monitor RSV release was consequently an important objective of this study. Using this assay, we identified drebrin as a cellular factor required for RSV release. We further showed that drebrin interacts directly *in vitro* and in cells with the viral matrix protein at viral budding sites. Analysis of the drebrin interactome during infection identified an additional cellular factor, TM9SF2, which is also required for RSV release and appears to contribute to the relocalization of drebrin to viral budding sites.

For many enveloped viruses, scission of the viral envelope from the plasma membrane is mediated by the ESCRT machinery. Viruses frequently recruit this pathway through their matrix proteins, which interact with specific ESCRT-III components (45, 46). In the case of RSV, a previous study reported that production of supernatant-associated virus was not affected by expression of a dominant-negative form of VPS4, the ATPase required for the final disassembly of ESCRT (26). Our results are consistent with this previous study as depletion of VPS4 did not impair virus release in the RSV release assay we established (Fig S5). In addition, we demonstrated that depletion of CHMP4B, a core component of ESCRT-III, also has no effect on viral release (Fig S5). Taken together, our results clearly support the conclusion that RSV release does not require the canonical ESCRT-III pathway.

Viral matrix proteins are likely to contribute directly to viral release, since direct or indirect interactions between matrix proteins and cellular scission factors are a recurrent feature, including viruses that use ESCRT-independent mechanisms. Our data strongly demonstrate that the cellular protein drebrin interacts with RSV M. This interaction was readily detected by co-immunoprecipitation from infected cells. Importantly, interaction assays performed using purified proteins demonstrated that the interaction is direct. Mapping experiments indicate that both the actin-binding domain (ABD) and intrinsically disordered domain (IDD) of drebrin are required for this interaction, since neither domain was sufficient when expressed separately. The requirement for the ABD may reflect a role for actin in stabilizing drebrin at the plasma membrane rather than a direct contribution of the ABD to M binding. Specifically, the ABD may anchor drebrin to actin filaments, thereby positioning the IDD domain for interaction with M. This interpretation is consistent with the observation that the IDD expressed alone was not efficiently targeted to filopodia. Nevertheless, we cannot exclude a direct contribution of the ABD to M binding and the surface of the ABD that is not engaged with actin could potentially interact with M. The corresponding region of M interacting with drebrin remains to be identified. Since M forms dimers and associates with the plasma membrane, drebrin may bind to a surface of the M dimer that is distinct from the region involved in membrane association. Defining the precise interaction sites within both proteins will be important for determining how the M-drebrin complex contributes to release.

Our results indicate that drebrin contributes to the release of RSV particles and suggests that the drebrin region required for M binding may be also required for its role in RSV release. Indeed, the ABDs-IDD mutant, which retained the ability to interact with M, was still able to increase the release of RSV, albeit to a much lesser extent than WT drebrin. This could be explained by the stronger interaction between this mutant and M, leading to excessive retention of M that may prevent the dynamic rearrangements required for productive particle release. Conversely, the ADFH-ABDs mutant, which did not interact detectably with M but remained associated with filopodia, was not only inactive but significantly reduced RSV release. This phenotype is consistent with a dominant-negative effect, potentially resulting from competition with endogenous drebrin for actin- or membrane-associated binding sites. Taken together, these results indicate that the drebrin domain required for RSV release could overlap with the domain involved in M interaction.

During infection, drebrin relocalized to the cell periphery and showed substantial colocalization with actin-rich membrane extensions. The presence of actin in M-drebrin immunoprecipitates further supports a close association between drebrin and actin cytoskeleton during infection. These observations are consistent with previously described roles of drebrin in membrane protrusion and filopodia formation (27), as well as the well-established direct interaction between drebrin and actin (36). In addition, we observed viral filaments emerging from filopodia enriched in both actin and drebrin. This finding is in agreement with previous descriptions of RSV-induced, Arp2/3-dependent, filopodia-like protrusions. As proposed in that study, actin-and drebrin-rich membrane extensions may contribute to the dissemination of RSV particles (20). The observation that the drebrin protein associates along RSV viral filaments by expansion microscopy suggests that the protein may be partially retained within viral particles, which would support a more specific role in membrane scission. Drebrin has additionally been implicated in ectocytosis, a process involving the release of extracellular vesicles from the plasma membrane that shares conceptual similarities with the release of enveloped viral particles. However, the molecular mechanism by which drebrin contributes to ectocytosis has not yet been established.

How might the M–drebrin interaction contribute to membrane scission or particle release? One possibility is that drebrin helps generate or stabilize the membrane curvature required for scission. A region within the drebrin helical domain could disengage from actin and insert into the membrane, thereby acting in a manner analogous to a BAR-domain-containing protein (47, 48). Many BAR domain proteins contain amphipathic helices and disordered regions that may contribute to membrane curvature. The presence of amphipathic helices in drebrin is not excluded and the IDD could also potentially contribute to such a function. Alternatively, drebrin may recruit or activate an as-yet unidentified ESCRT-independent scission machinery. A further possibility is that drebrin functions within a pathway involving Rab11-FIP2, a protein previously shown to promote the production of supernatant-associated RSV (26). Whether drebrin directly participates in membrane scission or instead serve as a scaffold for the recruitment of additional cellular fission pathway remains to be determined.

Characterization of the drebrin interactome during RSV infection identified TM9SF2 as a candidate required for efficient release of RSV particles. Moreover, depletion of TM9SF2 affected the relocalization of drebrin to virus-associated budding sites, suggesting that TM9SF2 may act upstream of drebrin recruitment. Interestingly, a recent study reported a role for TM9SF2 during the late stages of infection by influenza virus, and showed that TM9SF2 contributes to the maturation of the viral hemagglutinin protein (49). Further experiments will be necessary to determine whether TM9SF2 interacts directly with drebrin, with M, or with another component of the viral budding machinery, and whether it also regulates actin organization or membrane curvature.

In summary, this study identifies drebrin and TM9SF2 as cellular factors required for efficient RSV particle release and provides evidence for a direct interaction between drebrin and the RSV M protein. Our findings support a model in which infection-induced recruitment of drebrin to M-containing membrane domains, mediated at least in part by TM9SF2. Whether this pathway drives membrane curvature or directly catalyzes membrane scission remains to be established. Defining the molecular basis of this interaction should provide important insight into the poorly understood process of RSV release and may reveal a new mechanism of ESCRT-independent membrane scission.

## MATERIALS AND METHODS

### Cells

HEp-2 cells (ATCC, CCl-23) were maintained in Eagle’s minimum essential medium (MEM) supplemented with 10% heat-inactivated foetal bovine serum (FBS) and with 1% penicillin-streptomycin solution 5000 U/ml. The cells were grown at 37°C in 5% CO2.

### Viruses

The recombinant wild-type RSV and RSV-mCherry viruses are derivated from the human RSV Long strain (ATCC VR-26) and were previously described (9, 50). They were amplified on HEp-2 cells at 37°C (3 passages) and titrated by plaque assay (50).

### Plasmids

YFP-tagged DBN (drebrin-YFP) construct has been provided from addgene (#40359). Drebrin-YFP and YFP were PCR amplified and cloned into a modified form of the pCEP4 mammalian expression vector previously characterized in (51), between the NheI and NotI restriction sites. IDD-YFP truncation mutant was was kindly provided by Phillip R. Gordon-Weeks. The other truncation mutants (ADFH-ABD1-ABD2, ABD1-ABD2, ABD1-ABD2-IDD) were PCR amplified and cloned into the drebrin-YFP plasmid between the EcoRI and KpnI restriction sites. Finally, all the drebrin mutants were subcloned into the pCEP4 plasmid as the full-length drebrin-YFP. The pCI-M expression plasmid encoding the RSV-M protein was provided by Unité de Virologie et Immunologie Moléculaires, Université Paris-Saclay, France (18). The corresponding empty pCI vector (GenBank U47119) was used as a negative control. For expression and purification of recombinant M protein, a modified pCDF-Duet1 vector expressing a His_6_ fusion M protein, previously described in (42), was used. For expression and purification of recombinant P proteins, a codon optimised sequence of P was inserted in a pGEX4T-3 vector plasmid. pGEX-P_1-163_ and pGEX-P_127-241_ were generated using standard PCR, digestion, and ligation techniques. For expression and purification of recombinant drebrin protein, the pET24a-drebrin-His plasmid was used (addgene #40363).

### Cell infection

Viral stock was diluted in MEM or DMEM media devoid of FBS and then used to infect the cells at multiplicity of infection of 1. After 2 hours of adsorption, the inoculum was replaced by media supplemented with 2% FBS.

### Cell transfection

HEp-2 cells were transfected with the siRNAs listed in Table S2. Cells were seeded at 25,000 cells per well in 96-well plates and on 0,4 µm Transparent PET Membrane (Falcon; 353095) and at 125,000 cells seeded per well in 24-well plates. For the 96-well plates and 0.4 µm Transparent PET Membrane, the cells were transfected with siRNAs at a final concentration of 50 nM using 0.75 µL of Lipofectamine RNAiMAX (Thermo Fisher, cat. no. 13778150). For the 24-well plates, the cells were transfected with siRNAs at a final concentration of 30 nM with 1.5 µL of Lipofectamine RNAiMAX.

For transient plasmid transfection, 1 x 10^7^ HEp-2 cells were transfected with 16 µg of PCI-M plasmid DNA using Lipofectamine 2000 (Thermo Fisher, cat. no. 11668019) according to the manufacturer’s instructions. To generate HEp-2 cells stably harboring the replicating episomal pCEP4 plasmid, 1 x 10^7^ HEp-2 cells were transfected with 10 µg of pCEP4m plasmid DNA using Polyethylenimine (Polysciences; 24765). Two days post-transfection, the culture medium was replaced with MEM supplemented with 150 µg/mL hygromycin B gold (Invivogen; ant-hg-1). The selective medium was replaced every 2 days for 10 days to select cells containing the replicating episomal vector. The cells were subsequently maintained for 5 passages.

### Plaque titration assay

Forty-eight hours after transfection, siRNA transfected HEp-2 cells were infected with RSV at MOI of 1. At 24 h post-infection, the plates containing the infected cells were frozen at -80°C. Upon thawing, virus production was analyzed by plaque titration assay as previously described (53). Triplicates were realized per condition.

### Cell viability measure

Forty-eight hours after transfection, siRNA transfected HEp-2 cells were analyzed for the cell viability using the CellTiter-Glo kit (Promega, cat. no. G7571) and the Centro LB 960 luminometer (Berthold Technologies).

### Immunofluorescence

HEp-2 cells seeded and infected on coverslips were fixed with 4% PFA for 30 min and permeabilized for 10 min with 0,1% Triton-100X (Thermo Fisher; 10671652). Incubation with primary antibodies was performed for 1 h in PBS-BSA 1%. The primary antibodies used were as follows: drebrin (Progen; 612128 and abcam; 178408, 1/200), N RSV protein (INRAE; 1/40000), G RSV protein (Sigma; MAB858-2, 1/500) and M RSV protein (GeneTex; GTX640410; 1/100). Incubation with Alexa Fluor secondary antibodies (Thermo Fisher) and with phalloidin (Sigma; A22287, 1/800) was performed during 1 hour with Hoechst 33342 staining. Coverslips were mounted using Prolong Diamond Antifade Mountant (Thermo Fisher; 15468070). Images were captured by an Evident IXplore Spin SR Spinning Disk Confocal Microscope *with a 63x objective*. The entire volume of each cell was captured by performing a Z-stack acquisition (Zstep = 0.24µm). Images were analyzed using ImageJ software to quantify drebrin fluorescence signals. 3 Z steps were fused to perform the membrane quantification and drebrin staining was used to define the membrane. Briefly, the outer contour of each cell was manually delineated based on drebrin staining, generating a first set of Regions of Interest (ROIs). A custom macro was then applied to automatically generate a second set of ROIs corresponding to a region located 20 pixels inward from the outer contour, within the cytoplasm. This same macro was used to measure the mean fluorescence intensity within each ROI, yielding values for drebrin fluorescence intensity at the cell periphery and within the cytoplasmic compartment. The ratio of mean peripheral to cytoplasmic fluorescence intensities was then calculated for each cell to quantify drebrin relocalization to the cell periphery.

### Expansion microscopy

Cells fixed with PFA 4% were then permeabilizezd with and 0,1% Triton-100X (Thermo Fisher; 10671652). Incubation with drebrin primary antibody was performed for 1 h in PBS-BSA 0.1% and AF488 secondary antibody was added (Thermo Fisher, 1/100). Ultrastructured expansion protocol was conducted as previously described by Gambarotto et al. (54): cells were incubated overnight at 37°C in anchoring solution (Formaldehyde 0.7% / Acrylamide 1%). After gelation for 30 min at 37°C in monomer solution (Sodium Acrylate 23%, Acrylamide 10%, N,N’-methylenbisacrylamide 0.1% in PBS), gels were incubated in denaturation buffer (200 mM SDS, 200 mM NaCl and 50 mM Tris in ultrapure water) for 1.5 h at 95°C then rinsed twice in PBS. Incubation with anti-M primary antibody ((55) 1/500) then AF594 secondary antibody (Thermo Fisher, 1/200) and DAPI (1 µg/mL) were performed at 37°C for 3h and 2h respectively. Finally, the gel was expanded by incubating it in distillated water until reaching 4x expansion factor. Images were captured by an Evident IXplore Spin SR Spinning Disk Confocal Microscope *with a 63x objective*. The entire volume of each cell was captured by performing a Z-stack acquisition (Zstep = 0.8 µm). Images were analyzed using ImageJ software.

### Protein extraction, immunoprecipitation and Western blot

Total protein extracts were performed by adding 300µl of 1X NuPage LDS Sample Buffer (Invitrogen; NP0007) per 1 million cells and centrifugation with QIAshredder (Qiagen; 79654) was realized for the DNA lysis.

Immunoprecipitation experiments with infected extracts were performed as follows: twenty-four hours post infection cells were washed in PBS and lysis in the prepared buffer (PBS, protease inhibitor cocktail, 0,5% NP40) for 15 minutes at 37°C. Immunoprecipitations were performed by incubating indicated whole cell extracts overnight at 4°C with an GFP-Trap (Proteintech; 17383343). The beads were washed 4 times with a wash buffer (NaCl 150 mM, Tris-HCL 25 mM) then eluted in 1X NuPage. Immunoprecipitation experiments with transfected extracts were performed as follows: twenty-four hours post transfection cells were washed in PBS, cross-linked with Dithiobis Succinimidylpropionate (Thermo Fisher Scientific; 11821315) and lysis in the prepared buffer (PBS, protease inhibitor cocktail, 0,5% NP40) for 15 minutes at 37°C. Immunoprecipitations were performed by incubating the indicated whole-cell extracts overnight at 4°C with 2 µg of either an anti-drebrin antibody (Abcam, ab178408) or an anti-GFP antibody (Thermo Fisher Scientific, A11122), used as a negative control. Antibodies were pre-coupled to Protein A beads prior to incubation with the cell extracts. The beads were washed 4 times with a wash buffer (NaCl 150 mM, Tris-HCL 25 mM) then eluted in 1X NuPage LDS Sample Buffer.

Western Blot experiments were carried out as follows: protein extract was migrated in 4-12 % SDS-PAGE (Invitrogen; NW04125BOX) and transferred onto PVDF membranes (Invitrogen; IB24002), using the iBlot 2 Dry Blotting System, and incubated with corresponding antibodies in 5% milk. The primary antibodies used are the following: drebrin (Progen; 612128, 1/1000), RSV M (INRAE; 1/1000), GFP (Abcam; 11814460001, 1/2000), actin (Sigma; A2066, 1/1000), RSV (Abcam; Ab20745, 1/1000). Horseradish peroxidase-coupled secondary antibodies were obtained from Promega. Membranes were revealed by chemiluminescence (ThermoFisher Scientific, 34578). Densitometric quantification was performed on 3 independent experiments using Bio-Rad’s Image Lab software. The relative intensity of each protein band was measured and normalized to the corresponding loading control. For each condition, protein expression levels were expressed relative to the control condition. When no loading control was used, the relative band intensity of each condition was directly compared with the relative band intensity of the control condition.

### RT-qPCR

Cellular and viral mRNA were extracted using the RNeasy Mini Kit (Qiagen; 74106) and the QIAmp viral RNA extraction kit (Qiagen; 52904) respectively. Reverse transcription was performed with the reverse transcriptase M-MLV (Thermo Fisher; 28025013) and the qPCR with the FastStart Universal SYBR Green Master (ROX) (Roche; 4913914001). All experiments were realized according to the kit’s manufacturer’s instructions. Data were analyzed using the 2 ^- ΔΔCt^ method. GAPDH mRNA was used to normalize the mRNA quantity. Primer sequences are available on Table S3.

### RSV replication

Hep-2 cells were transfected with indicated siRNA and seeded in 96-well plates (0,25 × 10^5^ cells per well) in triplicate, 48h prior to infection with RSV-mCherry at a MOI of 1. mCherry fluorescence was monitored every 2h for 24h post-infection using an IncuCyte S3 live-cell imaging system (Sartorius), with images acquired using a 10X objective. Fluorescence signals were analyzed using the IncuCyte software by quantifying the total mCherry integrated intensity reported to the confluence (RCU × µm² per well per % of confluence) for each well at each time point. The percentage of the integrated mCherry intensity for each condition was calculated based on the average of the integrated mCherry intensity measured in the three wells at 24 hours after infection, relative to that of the siControl condition.

### RSV release assay

HEp-2 cells were seeded on 0,4 µm transparent PET Membrane (Falcon; 353095) and transfected or not with siRNA for 48 h. Cells were then infected with RSV-mCherry at MOI of 1. After 2 h of adsorption, “donor” cells were washed four times over 3 h with 2% MEM and subsequently placed in contact with “target” cells, seeded in a 24-well Permeable Support Companion Plate (Falcon; 353004) containing 2% OptiMEM for 20h. “Donor” cells were detached from the Transparent PET Membrane and transferred to a 24-well plate. The number of infected “donor” and “target” cells were quantified based on mCherry fluorescence using the Incucyte S3 system at 20h and 38h post-infection respectively (exposure time: 400ms; objective: 10X). At the end of image acquisition, a minimum cell size of 150 µm^2^ and a fluorescence threshold of 0.1-0.5 RCU were applied to all images. Raw data for “donor” cells, were obtained by selecting the Red Objects Count (ROC) parameter normalized to cell confluence, whereas raw data for “target” cells were expressed directly as ROC. Viral particle release efficiency was calculated as the ratio between the number of infected “target” cells (ROC) and infected “donor” cells (ROC normalized to percentage confluence). Finally, viral particle release efficiency for each condition was normalized to the corresponding control condition.

### Bacterial expression and purification of recombinant proteins

For drebrin expression, E. coli Rosetta 2 bacteria transformed with the pET24a-drebrin plasmid were grown from fresh starter cultures in Luria-Bertani (LB) broth at 37°C until reaching an OD_600_ of 0.8, followed by induction with 1 mM isopropyl-b-D-thiogalactopyranoside (IPTG) for 3 h at 28°C. Cells were harvested by centrifugation (4000 x g, 15 min) and then lysed in lysis buffer (50 mM Tris-HCl, 300 mM NaCl, 1 mg/mL lysozyme (Euromedex), 25 µg/ml RNase (Sigma) and protease inhibitors (Roche), pH 7.4) 1h on ice, followed by sonication (3 pulses of 15s at 50 mA, Qsonica Q500 sonicator). Lysates were clarified by centrifugation (15,500 x g, 30 min, 4°C), and the soluble His_6_-drebrin protein was purified on a nickel Sepharose column (HiTrap, 5 ml, IMAC HP; GE Healthcare). The bound protein was washed extensively with loading buffer (50 mM Tris-HCl, 300 mM NaCl, pH 7.4) plus 25 mM imidazole and eluted with a 25 to 500 mM imidazole gradient. 5mM dithiothreitol (DTT) and 2% glycerol were added to fraction containing His-drebrin, followed by concentration to 2 ml using Vivaspin20 columns (Sartorius Stedim Biotec). His-drebrin was then purified on a HiLoad 10/600 Superdex S200 column (GE Healthcare) in loading buffer plus 5mM DTT and 2% glycerol. The drebrin peak was concentrated to 3 mg/ml using Vivaspin20 columns. For M expression, E. coli Rosetta 2 bacteria transformed with the pCDF-M plasmid were grown in LB broth at 32°C until reaching an OD_600_ of 0.8, followed by induction with 0.5 mM IPTG for 4 h at 25°C. Cells were harvested by centrifugation and then lysed in lysis buffer (50 mM NaH_2_PO_4_-Na_2_HPO4, 300 mM NaCl, 1 mg/mL lysozyme, 25 µg/ml RNase, 0.2% CHAPS {3-[(3-cholamidopropyl)-dimethylammonio]- 1-propanesulfonate} and protease inhibitors, pH 7) 30 min on ice, followed by sonication. His_6_-M protein from lysates was purified on an IMAC colomn similarly to drebrin. The bound protein was washed extensively with loading buffer (50 mM NaH_2_PO_4_-Na_2_HPO4, 300 mM NaCl, pH 7) plus 25 mM imidazole and eluted with a 25 to 300 mM imidazole gradient. Fractions containing M were pooled, 1 mM DTT added and His tag cleaved with 2U/mL HRV 3C protease (His tagged, Sigma) while undergoing dialysis in a Spectra/Por porous membrane with a 12 to 14 kDa cut off (Spectrum labs) in dialysis buffer (loading buffer plus 1 mM DTT, pH 7) at 4°C overnight. Cleaved M protein was purified again on a nickel Sepharose column. Flow through containing M was concentrated to 2 ml using Vivaspin20 columns and purified on a HiLoad 10/600 Superdex S200 column in loading buffer. The M peak was exchanged to drebrin loading buffer and then concentrated to 3 mg/ml using Vivaspin20 columns. For P expression, E. coli BL21(DE3) bacteria transformed with pGEX-P-derived plasmids were grown at 37°C for 8 h in LB broth. Protein expression was induced by adding one volume of fresh LB medium with 0.4 mM IPTG for 16 h at 28°C. Bacterial pellets were resuspended in lysis buffer (20 mM Tris-HCl, 60mM NaCl, 1mM EDTA, 1 mg/ml lysozyme, 1 mM DTT, 0,1 % Triton X-100, pH 7.4) supplemented with complete protease inhibitor cocktail (Roche) for 1 h on ice. Benzonase (Millipore) was then added, and the lysate was incubated for 1 h at RT under rotation. The lysates were centrifuged at 4°C for 30 min at 15,500 x g. Glutathione-Sepharose 4B beads (GE Healthcare) were added to the clarified supernatants, and the mixtures were incubated overnight at 4°C under rotation. The beads were washed with lysis buffer, washed three times with phosphate-buffered saline (PBS), and then stored at 4°C in equal volumes of PBS. The glutathione transferase (GST) tag was removed by thrombin (Millipore) cleavage, and the cleaved product was exchanged into drebrin loading buffer. All P mutants were then concentrated and purified by size exclusion chromatography as described previously. The quality of protein samples was assessed by SDS-PAGE. Protein concentration was determined by measuring absorbance at 280 nm.

### Native gel

Protein samples were separated by electrophoresis on 1% agarose gel in Tris-Glycine buffer (10 mM Tris base, 100 mM Glycine). Recombinant proteins were incubated in Tris-NaCl buffer (50 mM Tris-HCl, 300 mM NaCl, 5 mM DTT, 2% Glycerol, pH 7.4) for 20 min at room temperature. Samples were mixed with 50% sucrose with bromophenol blue and run for 2 h at 80 V, following by staining with amido black.

### Sample preparation for mass spectrometry

Proteins affinity-purified with DBN-YFP, from RSV-infected and uninfected HEp-2 cells, were loaded on a SDS-PAGE (TGX precast gel 4-20%, Bio-Rad) and submitted to a short migration, in order to concentrate the proteins co-purified with DBN-YFP in one single gel band. For each sample condition, prepared in triplicate, protein bands were excised and subjected to in-gel tryptic digestion. Briefly, bands were washed with a solution of 30% acetonitrile in 70 mM ammonium bicarbonate with agitation at 800 RPM in a Thermomixer. Proteins were reduced with 10 mM dithiothreitol in 50 mM ammonium bicarbonate for 30 minutes at 56°C, followed by cysteine carbamidomethylation using 55 mM iodoacetamide in 50mM ammonium for 30 minutes at dark. After supernatant removal, the washing step was repeated. Finally, the gel bands were dried. Tryptic peptides were generated by addition of mass spectrometry-grade modified trypsin (Trypsin Gold, Promega) at 5 ng/µl in 50mM ammonium bicarbonate, and overnight incubation at 37°C at 400RPM. Tryptic peptides were extracted by addition of one volume of 50% acetonitrile in 0.1% formic acid and incubation during 20min at 800 RPM, before addition of one additional volume of 100% acetonitrile. The supernantant containing the tryptic peptides were vacuum-dried and resuspended in 2% acetonitrile in 0.05% trifluoroacetic acid for mass spectrometry analysis.

### Mass spectrometry analysis

NanoLC-MS/MS analysis were performed using a nanoElute liquid chromatography system coupled to a timsTOF Pro 2 mass spectrometer (Bruker, Billerica, MA, USA). Briefly, peptides were desalted online using a trap column (Waters, NanoE MZ Sym 18, 180 µm x 20 µm Trap column) before being loaded onto an Aurora analytical column (ION OPTIK, 25 cm x 75 µm, C18, 1.6 µm), and eluted with a gradient of 0–35% solvent B over 100 minutes as following: 0-15% B in 60min, 15-23% B in 30 min, 23-35% B in 10 min. Solvent A was 2% acetonitrile in 0.1% formic acid and water, and solvent B was 99.9% acetonitrile in 0.1% formic acid. MS and MS/MS spectra were recorded from m/z 100 to 1700 with a mobility scan range from 0.65 to 1.45 V_s/cm². MS/MS spectra were acquired using the PASEF (parallel accumulation-serial fragmentation) ion mobility-based acquisition mode, with the number of PASEF MS/MS scans set to 10.

### Mass spectrometry data analysis

Tandem mass spectra were processed and converted into mgf files using DataAnalysis software (Bruker, Billerica, MA, USA). Protein identifications were performed with the Mascot search engine (Matrix Science, London, UK; version 2.6.2) against the Uniprot human database (010322, 20,380 entries) including the DNB-YFP and trypsin protein sequences with a strict trypsin digestion, and up to 2 possible miscleavages. Peptides and fragments tolerances were set to 15 ppm and 0,05 Da respectively. Carbamidomethylation of cysteine was specified as a fixed modification and oxidation of methionines were specified as variable modifications. Label-free quantification was performed using spectral counting. Briefly, total spectral count values were extracted from Scaffold software (version scaffold 4.4.8, Proteome Software Inc, Portland, OR), filtered with a 95% probability and 0.1% FDR for protein and peptide thresholds, respectively. For statistical analysis, zero values present at the protein level were replaced with a constant value set at 0.1. Relative quantification between the two conditions was performed using the spectral count datasets and a beta-binomial test with the R package ’ibb’ (version 20.06). Only proteins with at least two spectral counts in at least two biological replicates of at least one condition were considered. Proteins were considered as significantly enriched in the DBN-YFP pull-down from RSV-infected compared to uninfected condition for p-value <0.05 and fold change greater than two.

## Statistical analysis

Statistical analyses are specified in each figure legend and were performed using GraphPad Prism 11.0.2 software. Each experiment was conducted using biological triplicates, unless otherwise indicated. Significance levels are indicated as follows: *\*\*\*\*: P* < 0.0001; *\*\*\*: P* < 0.001; *\*\*: P* < 0.01; *\*: P* < 0.05; *ns: P* > 0.05.

## ACKNOWLEDGMENT

We acknowledge the Proteomic-Gif (SICaPS) platform, supported by IBiSA, the Région Ïle-de- France (SESAME and DIM One Health programs), Plan Cancer, CNRS and the Paris-Saclay University. We thank Virginie Redeker for helpful discussion. We thank CYMAGES facility, part of UVSQ-Paris Saclay University. We acknowledge members of the team - Mécanismes moléculaires de la multiplication des Pneumovirus, for helpful discussions.

